# Biophysical modeling of excitation/inhibition imbalance in Alzheimer’s disease

**DOI:** 10.64898/2026.09.30.755800

**Authors:** Qin Deng, Cici Chen, Basilis Zikopoulos, Arash Yazdanbakhsh

## Abstract

**INTRODUCTION:** Alzheimer’s disease (AD) involves early sensory and perceptual changes that can precede memory decline. We developed a two-layer rate-based neural model of early and higher visual areas with feedforward, feedback, and recurrent connections to examine how excitatory/inhibitory (E/I) balance shapes visual motion perception in AD.

**METHODS:** Using Motion-Induced Position Shift and Altered Motion Repulsion stimuli, we factorially manipulated excitatory (E) and inhibitory (I) subfield power and connectivity to examine their effects on illusion representation across visual layers.

**RESULTS:** Increased E, decreased I, and increased E/I reduced illusion representation, consistent with reduced illusion susceptibility in AD, whereas the converse increased illusion representation, reflecting neurotypical processing. Higher-level layer 2 representations were more resistant to E/I perturbations than lower-level layer 1. Illusion representation was less affected by E/I increases in intra-areal (recurrent) connectivity than in inter-areal (feedforward/feedback) connectivity.

**DISCUSSION:** This mechanistic framework links E/I balance and connectivity to altered visual motion illusion representations in AD and suggests that E/I and connectivity measures may provide probes of early AD neurobiology. The framework may also generalize beyond vision.

## 1. Background

Alzheimer’s disease (AD) is a progressive neurodegenerative disorder characterized by cognitive decline, dementia, and eventual impairment in performing everyday activities. With increasing human life expectancy, the prevalence of AD has risen significantly, making it one of the most critical age-related health concerns of the 21st century, imposing immense burdens on healthcare systems and caregivers ^1–3^. Extensive research has centered on hallmark biomarkers of AD, such as amyloid plaques and neurofibrillary tangles, which are considered central to its pathology ^4–6^.

Recent studies have expanded the understanding of AD beyond amyloid and tau pathologies to include disruptions in broader neurobiological systems, such as the balance between excitatory and inhibitory (E/I) synaptic signaling, a critical indicator of healthy brain function ^7,8^. Increasing evidence suggests that disturbances in neural circuit function, particularly involving E/I imbalances, may play a role in disease progression.

The E/I balance is maintained by excitatory neurotransmission, primarily mediated by glutamate, and inhibitory neurotransmission, largely governed by GABA^9–12^. In AD, the excitatory components of neural circuits are vulnerable to pathological changes, manifesting as neural hyperexcitability, where excitatory inputs become excessively strong or are inadequately counterbalanced by inhibitory signals ^13,14^. Neural hyperexcitability has been directly linked to amyloid-beta plaque accumulation, which disrupts synaptic function by enhancing excitatory neurotransmission and impairing inhibitory control^13–15^. This imbalance aggravates synaptic dysfunction, neuronal death, and cognitive decline, contributing to abnormal neuronal firing patterns and impaired information processing in AD. Inhibitory mechanisms may also undergo pathological changes in AD, where GABAergic signaling and reductions in inhibitory interneuron activity further disrupt E/I balance ^11,16–19^. These changes impact brain networks and circuits stability, exacerbating hyperexcitability and contributing to cognitive and sensory processing deficits.

The pathological shift towards excitation in AD extends beyond cognitive deficits to affect sensory processing, including in the visual system. This system, characterized by a delicate interaction of excitatory and inhibitory signals, is highly susceptible to neurodegenerative changes. Functional disturbances in the visual system have been documented in AD patients, from lower-level deficits such as reduced visual acuity, color discrimination, contrast sensitivity, and visual processing speed, to higher-level deficits such as struggles in visual-spatial attention and the recognition of complex features like faces ^20–26^. These perceptual alterations may serve as markers of underlying neural dysfunction. For instance, the Motion-Induced Position Shift (MIPS) illusion, where the perceived alignment of moving Gabor patches distorts, is sensitive to E/I imbalances and has shown alterations in AD patients ^27^. Similarly, the Altered Motion Repulsion (AMR) illusion, where the perceived angular deviation between moving dots is reduced in AD patients, correlates with cognitive assessments like the Mini-Mental State Examination (MMSE) ^28^.

These visual processing changes allow exploration of the connection between E/I imbalances and sensory deficits in AD. By modeling these phenomena, we can investigate how specific neural circuit changes mimic sensory processing abnormalities in AD through reverse engineering and predict how altering E/I ratios impacts sensory perception through forward engineering.

In this study, we employed a spatio-temporal neural model to examine the effect of E/I imbalances on sensory processing, focusing on visual motion illusions such as MIPS and AMR. Our model simulated lower and higher-order cortices of the human visual system, including parameters like receptive field on-center/off-surround properties, recurrence strength (local intra-areal connectivity), feedforward, and feedback projections from higher visual areas ^29–36^. By adjusting parameters of these circuits and network to replicate E/I shifts observed in AD, we aimed to quantify the degree of visual system aberrations of perceptual representation.

This work connected known E/I imbalances in AD with observed sensory processing deficits to infer and advance our knowledge on neural circuit alterations reflected in sensory dysfunction. The findings have the potential to support the development of novel diagnostic tools and therapeutic strategies focusing on E/I balance and sensory deficits in AD, with the hope of timely management and treatment of the disease.

## 2. Methods

### 2.1 Neural model

This study implemented a rate based neural model, consisting of excitatory (E) and inhibitory (I) components, possessing on-center off-surround receptive fields. The model’s on-center off-surround connectivity within (recurrent) and between model brain areas featured excitatory neurons forming direct synapses with the postsynaptic neurons. In vivo, inhibition occurs indirectly via inhibitory interneurons synapsing onto postsynaptic neurons; however, the model simplified this by implementing direct monosynaptic inhibition, bypassing the indirect excitation of presynaptic inhibitory neurons. Our study adopted the on-center off-surround connectivity pattern, where the on-center reflected the E presynaptic contribution and the off-surround represented the I surround suppression mediated by interneurons. As shown in Figure 1A, we simulated the on-center off-surround recurrent/interareal connectivity using the Difference of Gaussians (DoG), implemented as the weighted element-wise subtraction of the inhibitory Gaussian (purple curve) from the excitatory Gaussian (blue curve) ^35,36^. To quantify the strength of E and I connectivity, we used two parameters: the projection extent (width) of E and I connectivity, denoted by *σ_ext_* and *σ_in_*_ℎ_, and the projection strength (amplitude) of E and I connectivity, denoted by *α_ext_* and *α_in_*_ℎ_. Together, *σ* and *α* quantify the overall effective E and I *power*, to be used for assessing E/I balance.

**Figure 1.**
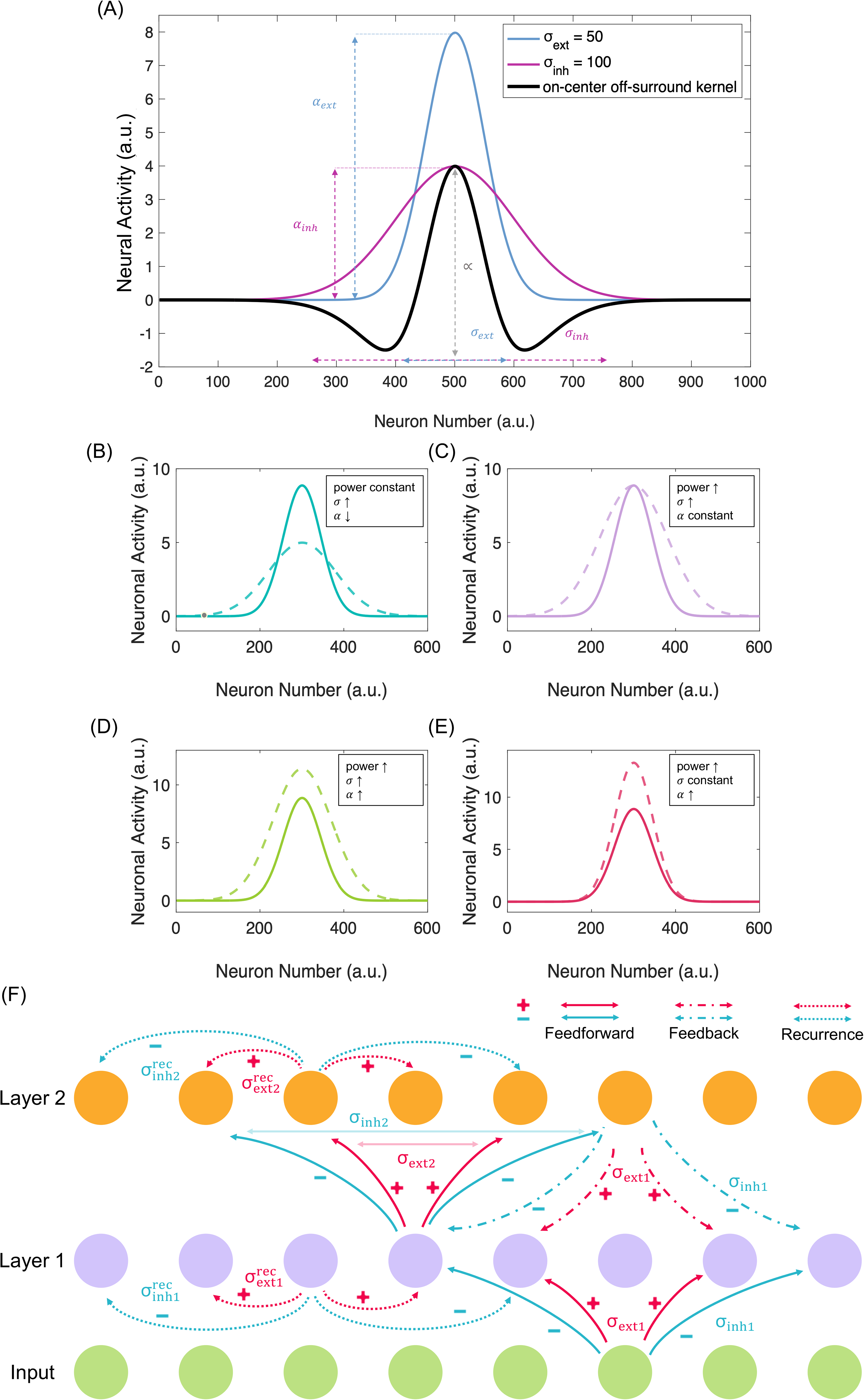
The neural model feedforward, recurrent, and feedback connectivity and architecture. (A) An example of the neural model on-center off-surround (black line) with *σ* as the projection extent (width) and *α* as the strength of projection (amplitude). Connectivity weights from the middle neuron (500) to the 1000 model neurons are shown; neuronal population induced activation from neuron 500 is proportional to respective connection weights. *σ* and *α* together quantify the overall E and I power. Excitatory component with *σ_ext_* and *α_ext_* is represented in blue gaussian and inhibitory component with *σ_in_*_ℎ_ and *α_in_*_ℎ_ is represented in purple gaussian. The difference of the two gaussians (DoG) representing the on-center off-surround model is depicted in black. Subplots (B-E) show four possibilities of increased excitation (E) or inhibition (I): increased *σ* can represent an expansion in the width of E (or I) spread while increased *α* (amplitude) can represent an enhancement of E (or I) strength. (B) Increasing *σ* to expand E (or I) while retaining the total E (or I) power represented by area under the curve, resulting in concurrently decreased *α*, (C) increasing *σ* with a maintenance of *α*, thus increasing the overall E (or I) power represented by increased area under the curve, (D) simultaneously increasing *σ* and *α*, thus increasing the overall E (or I) power represented by increased area under the curve, (E) increasing α with a maintenance of *σ*, thus increasing the E (or I) represented by increased overall area under the curve. (F) The proposed neural model with input layer, layer1, and layer2, with neurons connected by feedforward, feedback, and recurrent E and I connectivities. The extent of connectivity is represented by *σ*_*ext*1_, 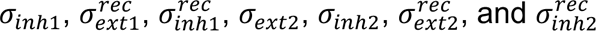.

In addition, there were several ways to achieve an overall increase in E (or I) power (Figure 1B–E): First, by increasing *σ* of the Gaussian, *α* decreased (constant power), an intrinsic characteristic of normal distribution Gaussian curve (Figure 1B). Increasing *σ*, and maintaining peak of the Gaussian, increased the area under the curve (power) (Figure 1C). Scaling up the curve in 1C made both α and *σ* larger, thus further increasing the overall area under the Gaussian, thus power (Figure 1D). Alternatively, only increasing *α* while keeping *σ* constant, increased power (Figure 1E). Illustrations in Figures 1C-E were the biologically plausible cases that we considered for the results section.

The input first stimulated the layer 1 neurons presented in purple via an on-center off-surround pattern (Figure 1F). Layer 1 represented lower visual processing levels such as the retina, LGN, and V1. The information flowing from layer 1 to layer 2 represented a feedforward connectivity pattern that consisted of both E and I components. Layer 2 represented higher visual processing levels in the hierarchy such as V2, V3, V4, and Medial Temporal area (MT). In our study, the receptive field ratio of layer 1 to layer 2 was 1:2, following physiological data from primates ^37^. The information flowing from layer 2 to layer 1 represented a feedback connectivity pattern with *σ_ext_* and *σ_in_*_ℎ_. The recurrence (intra-areal connectivity) of layer 1 and 2 was simulated by within-layer on-center off-surround connectivity.

To adapt the model for the computational approach, the integrated spatiotemporal activity in the visual area was modeled based on the Euler method:

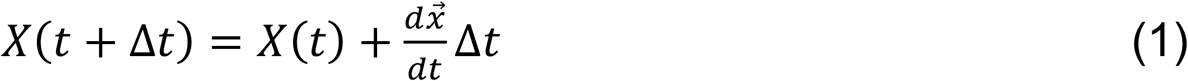

The Euler method splits time into many time intervals so that the neural activity changes over time can be quantified within each time interval. In each time interval, the spatiotemporal equation was used. The spatiotemporal equation describes a rate based neural model ^30–35,38^:

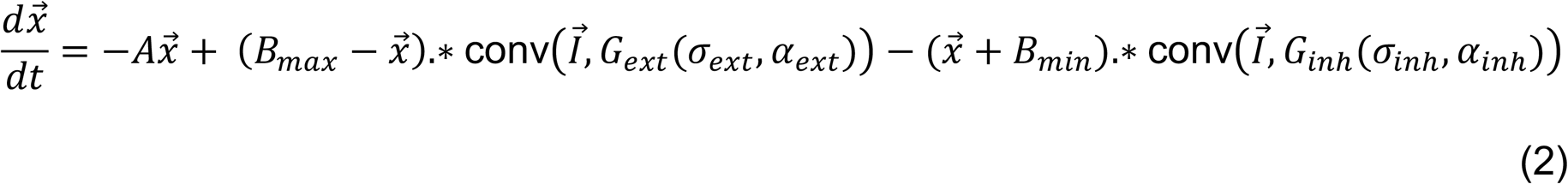

Vector *x⃗* was the spatiotemporal response of a particular model layer, each layer composed of activities of x number of neurons. *dx⃗*/*dt* represented the evolving activity of neurons in space over time. *A* was the decay rate that returned the model neuron activity to a resting state when there was no presynaptic input. *B_max_* was the maximum activity level of neurons whereas *B_min_* was the minimum activity level of neurons. For MIPS and AMR, the model had a decay rate of *A* = 0.5 and *A* = 10 respectively, maximum activity level *B_max_* = 90, and minimum activity level *B_min_* = 10. By using the Euler method, we approximated neural activity at each time step considering Equation 2. *G_ext_* was the excitatory Gaussian with an excitatory on-center width of *σ_ext_* and amplitude of *α_ext_*; *G_in_*_ℎ_ was the inhibitory Gaussian with an inhibitory off-surround width of *σ_in_*_ℎ_ and amplitude of *α_in_*_ℎ_. The convolution operation applied the filter-like Gaussian to the input vector *I*, in which the modification of *σ* and *α* altered the power of the E or I neural projection.

By considering the range for *σ_ext_*, *σ_in_*_ℎ_, *α_ext_*, *α_in_*_ℎ_, *B_max_*, and *B_min_* parameters of the model for neurotypical and Alzheimer disease conditions, based on (*σ_ext_* × *α_ext_* × *B_max_*)/(*σ_in_*_ℎ_ × *α_in_*_ℎ_ × *B_in_*_ℎ_), we estimated our model power ratio of E to I to be 0.4:1 to 23:1, which has overlap with the reported physiological range of 3:1 to 9:1, thus showing we had exhaustively conducted a parameter search within and beyond the physiological range^39^. The *σ_ext_* to *σ_in_*_ℎ_ was chosen with a 1:2 ratio, aligning with the past modeling practices ^40^ (Ch3.3 pg.136 1:1.5, Ch2.2 pg 62 1:1.6 ext:inh). We incorporated this 1:2 ratio of *σ_ext_* to *σ_in_*_ℎ_ when calculating the effective power ratio of E to I. Moreover, anatomical findings suggest that there are 70-80% of excitatory pyramidal neurons and 20-30% of inhibitory interneurons in the neocortex ^41–43^. These inhibitory interneurons are capable of receiving stimuli from both excitatory and inhibitory synapses, which could either account for more E or more I, creating a broader potential range of effective power ratio between E and I ^44,45^, and therefore justify our more exhaustive parameter search.

We further utilized the spatiotemporal model to simulate the within-layer recurrence by on-center off-surround communication within each of the two model layers: both layer 1 and layer 2 recurrently input themselves besides interlayer input, for example, layer 1 received the stimulus, within-layer recurrence, and feedback from layer 2. The overall input to layer 1, combining illusion input *I_input_*, recurrence *R_layer_*_1_, and feedback from layer 2 *x_layer_*_2_ was defined with the equation:

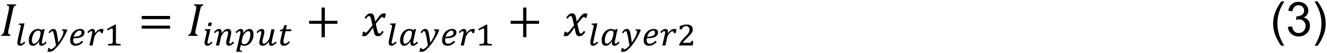

where *I_input_* was the initial illusion stimulus, *x_layer_*_1_ was the recurrence, and *x_layer_*_2_ was feedback from layer 2. Meanwhile, layer 2 received feedforward input from layer 1 *x_layer_*_1_ and within-layer recurrence *x_layer_*_2_:

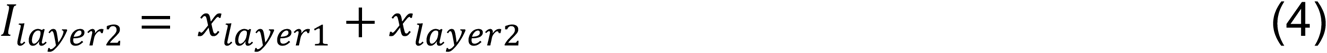

### 2.2. Motion-induced position shift

Motion-induced position shift (MIPS) is a visual illusion where the observers see the position of an object being displaced along the direction of motion. MIPS is affected in the early stage of AD when patients become less sensitive to the illusion ^27,46^. Our neural model for MIPS considered feedforward, feedback, and recurrent neural connectivity. Therefore, based on the changes in the neural connectivity and the implementation of these changes in the model, we could quantify the reduced illusion representation. Moreover, we could explore additional neural connectivity changes beyond the existing findings in AD that would result in reduced illusion representation and further propose such changes as candidates of underlying neural changes in AD or other neurodegenerative diseases.

Two vertically aligned Gabor patches were used as the input (Figure 2A). The outlines of the two Gabors were static, but the top had a rightward-shifting internal waveform carrier while the bottom had a leftward-shifting internal waveform carrier (Figure 2A–B) (Eq.5). We used the following sine function for the internal waveform carrier input:

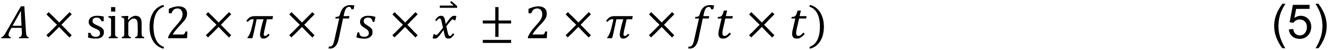

**Figure 2.**
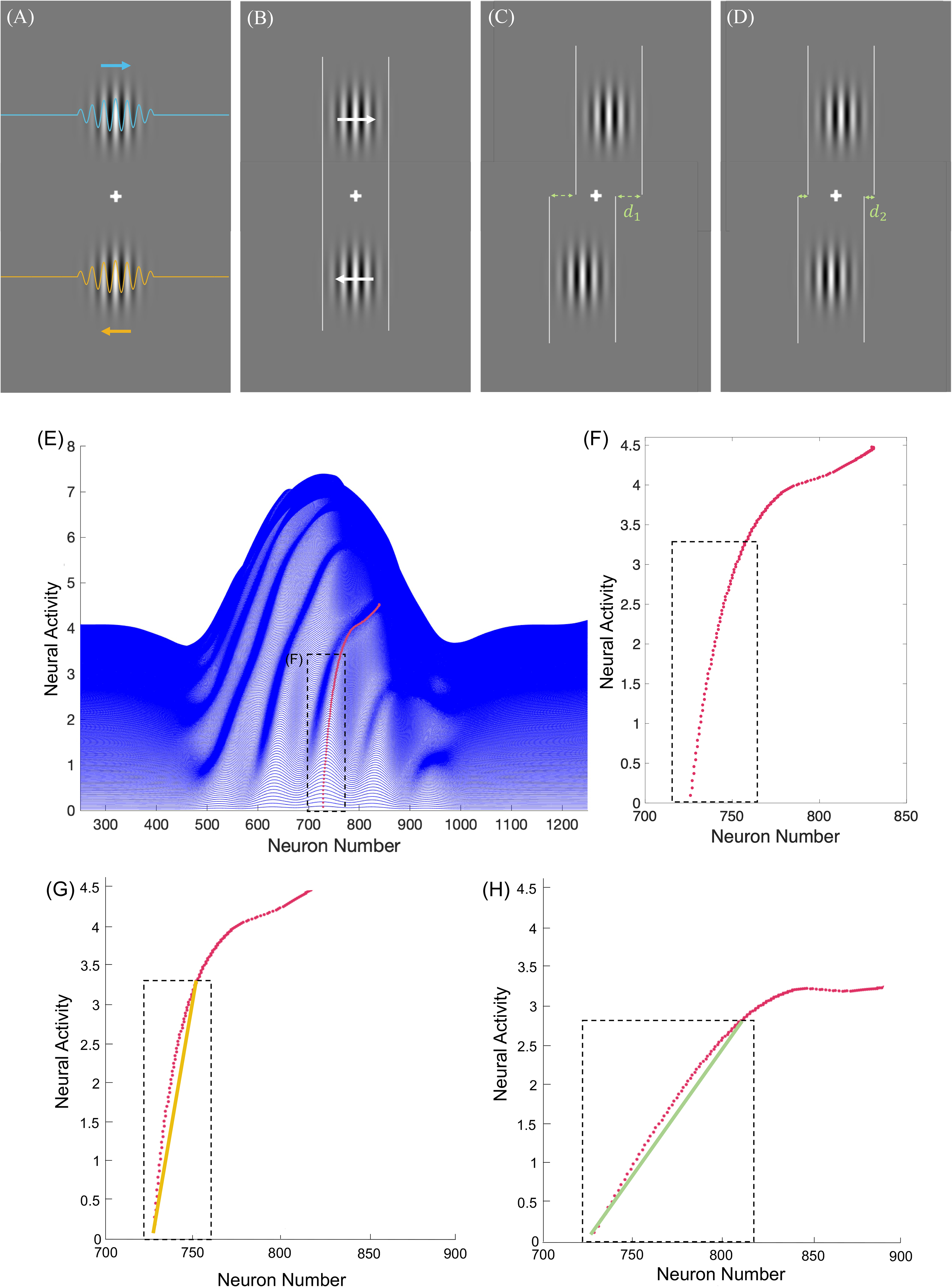
Motion-Induced Position Shift (MIPS) illusion input and its representation by the neural model. (A) Displays the rightward (blue) and leftward-moving (orange) sine functions as the sinusoidal base of the shifting internal waveform carrier. (B) Captures a snapshot of two vertically aligned Gabor patches with static outlines as inputs to the neural model at a point in time. The top grating has a rightward-shifting internal waveform carrier while the bottom has leftward-shifting internal waveform carrier. (C) Represents the illusion perception of neurotypical subjects, where the Gabor patches are perceived to offset significantly with a misalignment distance of *d*_1_. (D) Represents the illusion perception of individuals with AD, where the misalignment distance, *d*_2_, is 16 arcmin less than *d*_1_. Subplots (E) and (F) show Motion Induced Position Shift neural response of a right moving Gabor patch and its quantification. (E) The tilted trace of the local maxima (red dots) demonstrates the rightward shift induced by the motion. (F) 30 points (maximum neuronal activities over time, black box) were used to calculate the MIPS displacement representation; number 30 is an example representing the initial transient neuronal activity change related to motion encoding. Subplots (G) and (H) show a schematic diagram of MIPS quantification. Model neuron array activity was recorded after presenting the model with MIPS inputs; peaks of activity are illustrated by red dots. To quantify the strength of MIPS illusion representation, 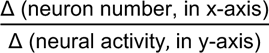 was calculated within the black box containing the red dots each at the peak neural activity as in (E), which measured the displacement of the neuron position with the peak activity, normalized by value change of the peak neural activity as an average of the represented position shift. In (G), the illusion representation is less than (H) since the same amount of activity spans across a smaller range of neurons.

in which *ft* = temporal frequency, *t* = time, *fs* = spatial frequency, and *x⃗* = position. A negative temporal component (−2 × *π* × *ft* × *t*) indicated a rightward-shifting internal waveform carrier whereas the positive one (+2 × *π* × *ft* × *t*) indicated a leftward-shifting internal waveform carrier. The alternating local peaks and troughs represented the alternating white and black regions of the Gabor patches (Figure 2A).

Furthermore, the Gabor patches had a decreasing intensity of black-white contrast from the center to the periphery of the Gabors. The Gabor patch outline was where stimulus contrast was low enough to perceptually blend with the gray background. Such contrast modulation leading to these peripheral low-contrast regions would make the overall illusion more apparent ^47^. To generate a Gabor input with an outline of low stimulus contrast, we point-by-point multiplied the sine function (Eq.5) with a Gaussian, which was then input into the two-layered MIPS neural model.

In neurotypical subjects, the Gabors are perceived to offset significantly, each displaced in the direction of the shifting internal waveform carrier (Figure 2C). In AD patients, the offset is largely reduced (Figure 2D). In AD subjects, the misalignment distance, *d*, is approximately 16 arcmins less than that of neurotypical subjects ^27^.

The model contained a range of *σ* and α parameter values as well as the combined power range values, *σ* × *α* (Table 1). In column 2 of Table 1, there are 8 *σ* parameters. Each *σ* corresponds to an *α* parameter in column 4 to achieve the representation requirement of Figure 1D. Each *σ* parameter and its corresponding α parameter has a range of 80 values, shown in columns 3 and 5 respectively. Thus, we ran 320 (4×80) permutations for each layer, with a total of 640 permutations from two layers. For each simulation run, we altered a single *σ* parameter (e.g. *σ_ext_*_1_ from 11 to 90), along with the associated α (e.g. α_ext1 from 1.01 to 1.8), while keeping other *σ* and *α* parameters constant (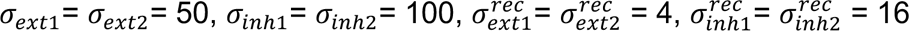, α=1.01), resulting in a specific *σ* × *α* power value and a two-dimensional model illusion representation graph. For example, one simulation run had the following parameters: 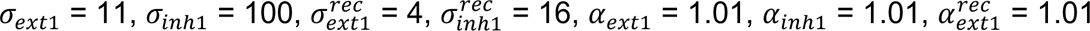, 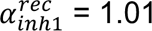; another simulation run had the following parameters: *σ_ext_*_1_ = 12 and *α_ext_*_1_ = 1.02 while other parameters remained the same. Then, we also performed parameter search where both *σ_ext_* and *σ_in_*_ℎ_ were variables, altered along with their associated *α_ext_* and *α_in_*_ℎ_, resulting in a three-dimensional model illusion representation space. For example, one simulation run had the following parameters: 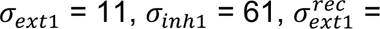 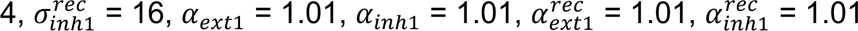; run had the following parameters: *σ_ext_*_1_ = 12 and *α_ext_* = 1.02 while other parameters remained the same.

**Table 1:**
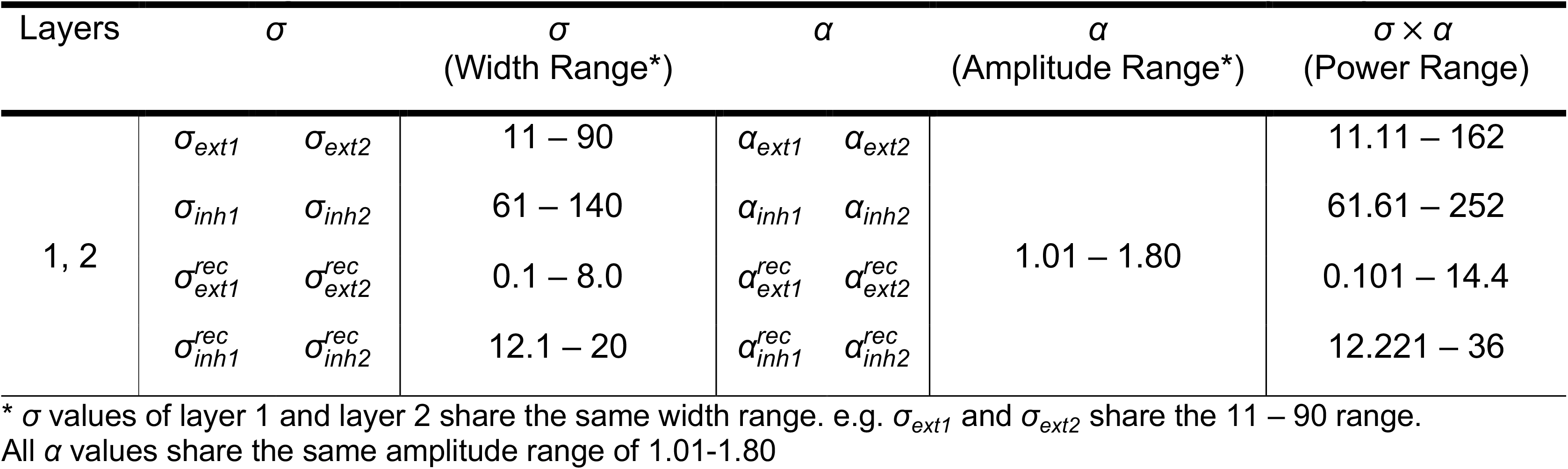
Range of parameter values in Motion-Induced Position Shift (MIPS)

| Layers | $\sigma$ | | $\sigma$<br>(Width Range*) | $\alpha$ | | $\alpha$<br>(Amplitude Range*) | $\sigma \times \alpha$<br>(Power Range) |
| --- | --- | --- | --- | --- | --- | --- | --- |
| 1, 2 | $\sigma_{ext1}$ | $\sigma_{ext2}$ | 11 – 90 | $\alpha_{ext1}$ | $\alpha_{ext2}$ | 1.01 – 1.80 | 11.11 – 162 |
| | $\sigma_{inh1}$ | $\sigma_{inh2}$ | 61 – 140 | $\alpha_{inh1}$ | $\alpha_{inh2}$ | | 61.61 – 252 |
| | $\sigma_{ext1}^{rec}$ | $\sigma_{ext2}^{rec}$ | 0.1 – 8.0 | $\alpha_{ext1}^{rec}$ | $\alpha_{ext2}^{rec}$ | | 0.101 – 14.4 |
| | $\sigma_{inh1}^{rec}$ | $\sigma_{inh2}^{rec}$ | 12.1 – 20 | $\alpha_{inh1}^{rec}$ | $\alpha_{inh2}^{rec}$ | | 12.221 – 36 |

\* $\sigma$ values of layer 1 and layer 2 share the same width range. e.g. $\sigma_{ext1}$ and $\sigma_{ext2}$ share the 11 – 90 range. All $\alpha$ values share the same amplitude range of 1.01-1.80

To quantify the model MIPS illusion representation, we found it best to capture the maximal transient part of the model neural activity because motion perception relies on the transient nature of inputs over time and temporal neural encoding. First, we found the local maxima of neural activity at each timestep and traced these peaks consecutively over 449 ms (Figure 2E) (red dots); each timestep neural activity had been shown with a plot, the next timestep plot was stacked on top, because continued input exposure increased the neurons’ activity. Then, we considered the peak that started at position 726 at time *t* = 1 and persisted in all the parameter scopes in Table 1. The peak traces (Figure 2F) formed a steep slope initially, representing a high neural activity buildup, then the slope gradually decreased over time. We calculated the initial slope to determine a cut-off point of high neural activity. Then, we found the critical point where the slope was 85% of the initial slope (15% reduction). This value was based on using 85% correct performance within psychometric functions for analytical purposes such as estimating just noticeable differences (JND) ^48,49^. All traced peaks from the initial point to the critical point were defined as the transient region of the neural activity registration. We then selected the first 60 peak points within the transient region to apply to all simulation runs (boxed region in Figure 2E–F). Finally, to quantify the model representation of visual misalignment distance d (Figure 2D), we used the following equation:

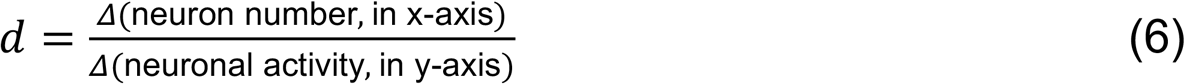

Where x represented the neuron position and y represented the strength of the neural activity at that position. The equation measured the overall increments of neuron position normalized by the value change of the peak neuronal activity to represent an average position shift. Figure 2G shows less illusion representation with a relatively small change in neuronal position (ΔX) and a relatively large change in neuronal activity (ΔY), resulting in an overall small ΔX/ΔY, suggesting a more limited span of activity, hence representing a weak illusion representation, such as that seen in AD individuals. In comparison, Figure 2H shows a relatively large ΔX with a similar amount of ΔY, resulting in an overall larger ΔX/ΔY, suggesting a more extensive span of activity hence representing a strong illusion representation, such as that seen in neurotypical individuals (Eq.6). For practicality, we kept ΔX as 8 based on how we conducted the parameter search. In summary, we considered this quantification a plausible representation of displacement because motion registration is based on transient stimuli and transient underlying neural activity. By choosing the boxed region in Figure 2E-F, we were picking the initial and transient model response to the stimulus.

### 2.3 Altered Motion Repulsion

We concurrently explored the model representation of another illusion, the Altered Motion Repulsion (AMR). In AMR, the angular separation between the two directions of motion is altered (perceived to be augmented) compared to its true value in neurotypical subjects. However, in AD, this alteration is reduced. The degree of reduced augmentation in angular separation correlates with the Mini-Mental State Examination (MMSE), a cognitive assessment for dementia and AD in the elderly ^28^. We investigated which parameters of feedforward, feedback, and recurrence of neural connections could reduce the amount of model representation of angular repulsion, which parallels AD patient’s reduced perceived augmentation in angular separation. With this method, we can propose a list of individual or combined feedforward, feedback, and recurrent factors as candidates for underlying changes in AD and other neurodegenerative disorders.

In order to investigate the model representation of AMR, which is originally based on Random Dot Kinematograms (RDK) in human subject experiments, we used a simplified version of the RDK as the input, i.e. two dots in motion which intersect each other as candidate interweaving and crossing RDK dots, each dot representing one of the two groups of RDK moving dots. One dot moves in the reference direction, which is 0 degrees to the right (horizontally); the other dot moves in the test direction, which deviates from the reference direction by an angle. This angle, known as the actual angular separation, has the following range of values: 0°, 5.6°, 11.2°, 22.5°, 45°, 67.5°, and 90° ^28^, which we also used as input to the model. To eliminate the direction aliasing effect due to discrete pixel input organization in the model, we rotated the reference and test dot trajectories so that the two trajectories are symmetric with respect to a horizontal line (Figure 3A).

**Figure 3.**
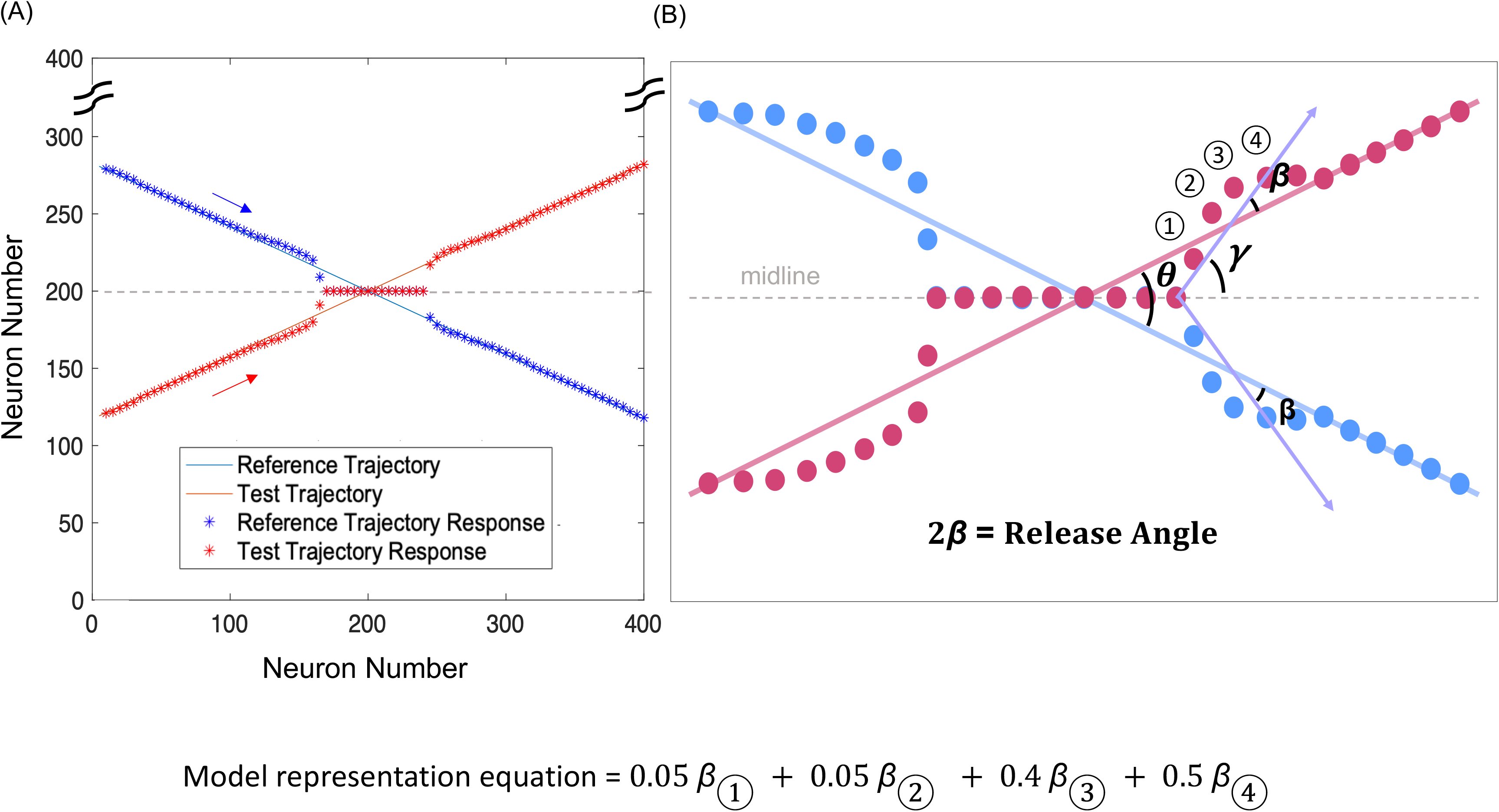
Altered Motion Repulsion (AMR) model input, representation, and its quantification. (A) The blue line shows the reference trajectory with the blue arrow indicating the direction; the red line shows the test trajectory with the red arrow indicating the direction. The true values of the angular separation are 0, 5.6, 11.2, 22.5, 45 (shown in this figure), 67.5, and 90. The blue asterisks show the maximum neural activity at various locations along the reference trajectory, whereas the red asterisks show the maximum neural activity at various locations along the test trajectory. (B) The model representation of angular repulsion was quantified by the magnitude of the release angle (deviation from true angle), 2*β* = 2(Q − N/2). Peak neuronal activities deviating from true motion trajectory (release points) ①, ②, ③ and ④ are shown. Model representation equation = 0.05 *β*_①_ + 0.05 *β*_②_ + 0.4 *β*_③_ + 0.5 *β*_④_.

To set the reference and test dot trajectories, we implemented polar coordinates system to translate the true angular separation into x, y values:

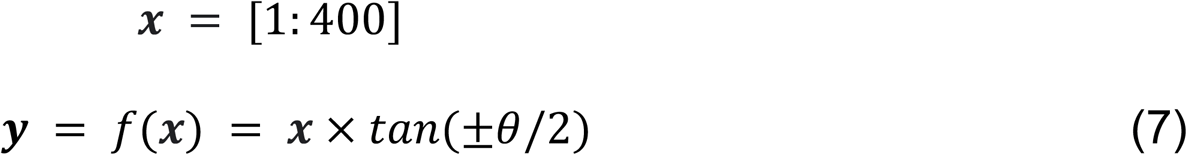

*x* was an array of the horizontal positions of both the reference and test dot trajectories, *y* was an array of the vertical positions as a function of *x*, and O was the actual angular separation between test and reference trajectories. −N/2 represented the angle between the midline and the reference dot trajectory whereas +N/2 represented the angle between midline and the test dot trajectory.

To investigate the impact of individual *σ* and α parameters in AMR, we first incorporated a range of *σ* and *α* values, as shown in top part of Table 2. Unlike in MIPS, where *σ* had a corresponding *α*, in AMR, both *σ* and *α* served as independent variables. Next, to explore the effects of E and I power, as well as changes in the E/I ratio in AMR, we established a baseline as a control condition using a specific set of *σ* and *α* parameters (Table 2, shaded row). To determine how different E/I ratios influence the degree of angular repulsion, we examined two main scenarios: an increase in the E/I ratio and a decrease in the E/I ratio relative to the baseline. Then, we presented 10 different combinations of changes in individual E and I components, which contributed to either an increased or decreased overall E/I ratio (lower part Table 2). Since the E/I ratio depends on both *σ* and *α* parameters (Figure 1C–E), we found respective *σ* and *α* values for each of the 10 combinations. The corresponding E and I power values were derived by multiplying the respective *σ* and *α* parameters. The E/I ratio was calculated by dividing E power by I power.

**Table 2:**
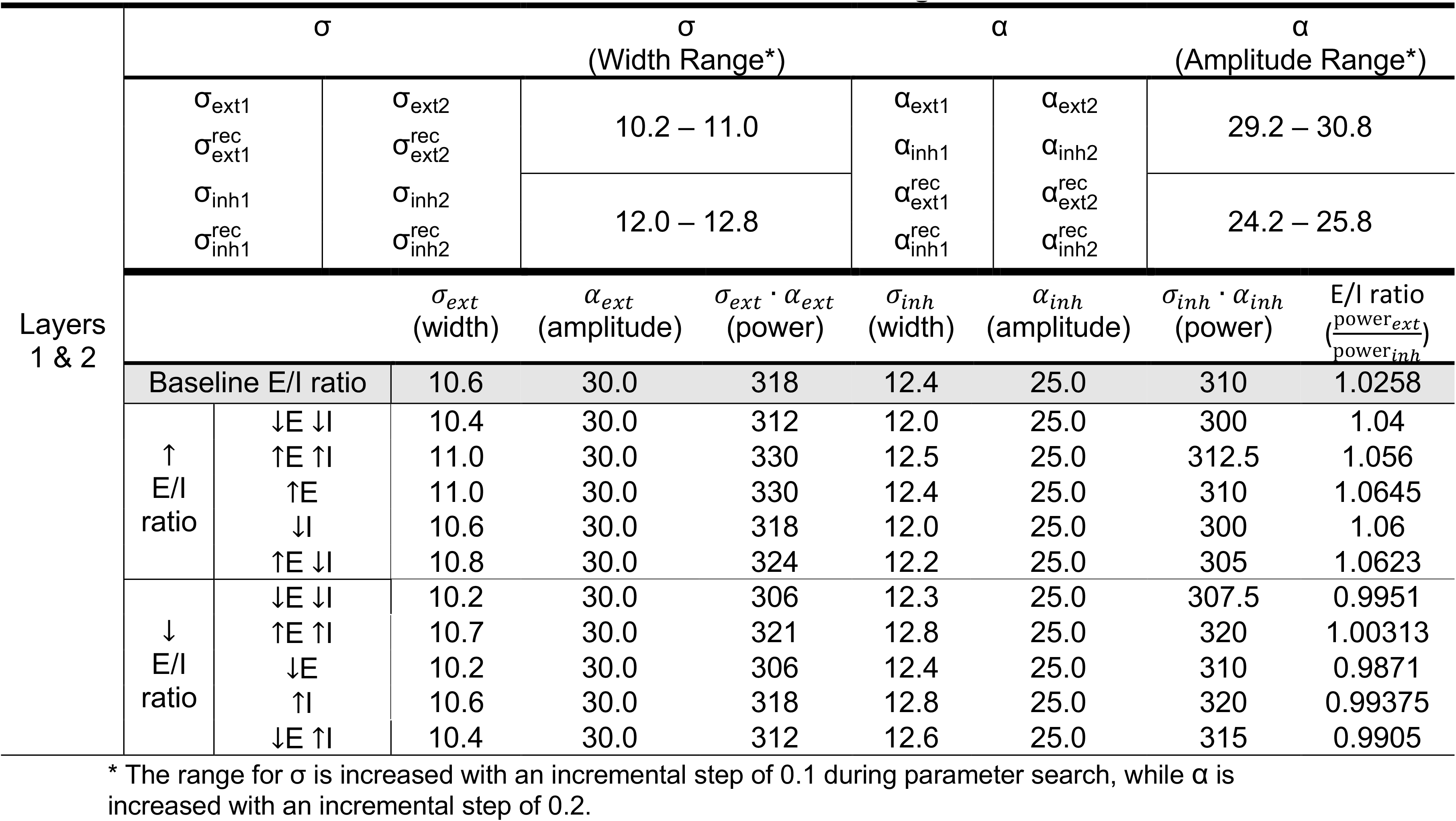
Range of parameter values in Altered Motion Repulsion (AMR) and 10 combinations of E and I subfield changes for AMR.

To quantify the amount of angular repulsion, we defined the release angle (2*β*) as the angle between the actual (N) and augmented (Q) angular separation:

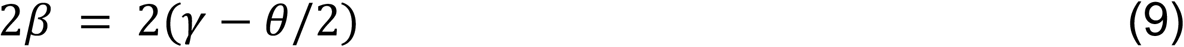

Finally, we calculated the weighted average of the release angle 2*β* of the first four release points for the baseline condition to reflect the distributed read out of neuronal activity in the vicinity of moving dots intersection, 5 E/I ratio increase possibilities, and 5 E/I ratio decrease possibilities for comparison of the amount of angular repulsion.

## 3. Results

### 3.1 The Effects of E and I Subfield Power in Altering Motion-Induced Position Shift (MIPS)

In order to understand the underlying neural mechanisms behind changes in susceptibility to the Motion-Induced Position Shift illusion, we examined the strength of model illusion representation when excitatory (E) and inhibitory (I) subfield power in first- and second-layer feedforward, feedback, as well as recurrent connectivity were modified. For more definitions of subfield power, refer to the methodology section (Figure 1B–E). The results of different modifications in subfield powers are shown in Figure 4. Overall, the results showed that increases in E subfield power in the first and second layers, as well as their recurrent connectivity, corresponded to reduced representation of the illusion. On the other hand, when I subfield power in the first and second layers, as well as their recurrent connectivity increased, the amount of illusion representation increased (Figure 4). These two major themes in E and I subfield power changes were further confirmed by the significant positive or negative Kendall, Pearson, and Spearman correlation coefficients denoted by *τ*, *r*, and *ρ* respectively (Supplementary Table 1). The rank-based Kendall *τ* and Spearman *ρ* correlation had -1 for all E subfield power modification and 1 for all I subfield power modification in relation to the MIPS model illusion representation, showing pure monotonic increasing and decreasing trends respectively. Similarly, the absolute values of the parametric Pearson r correlation coefficients were all above 0.8, suggesting significant parametric correlations between the E and I subfields power and the model illusion representation (p-values close to 0, Supplementary Table 1).

**Figure 4.**
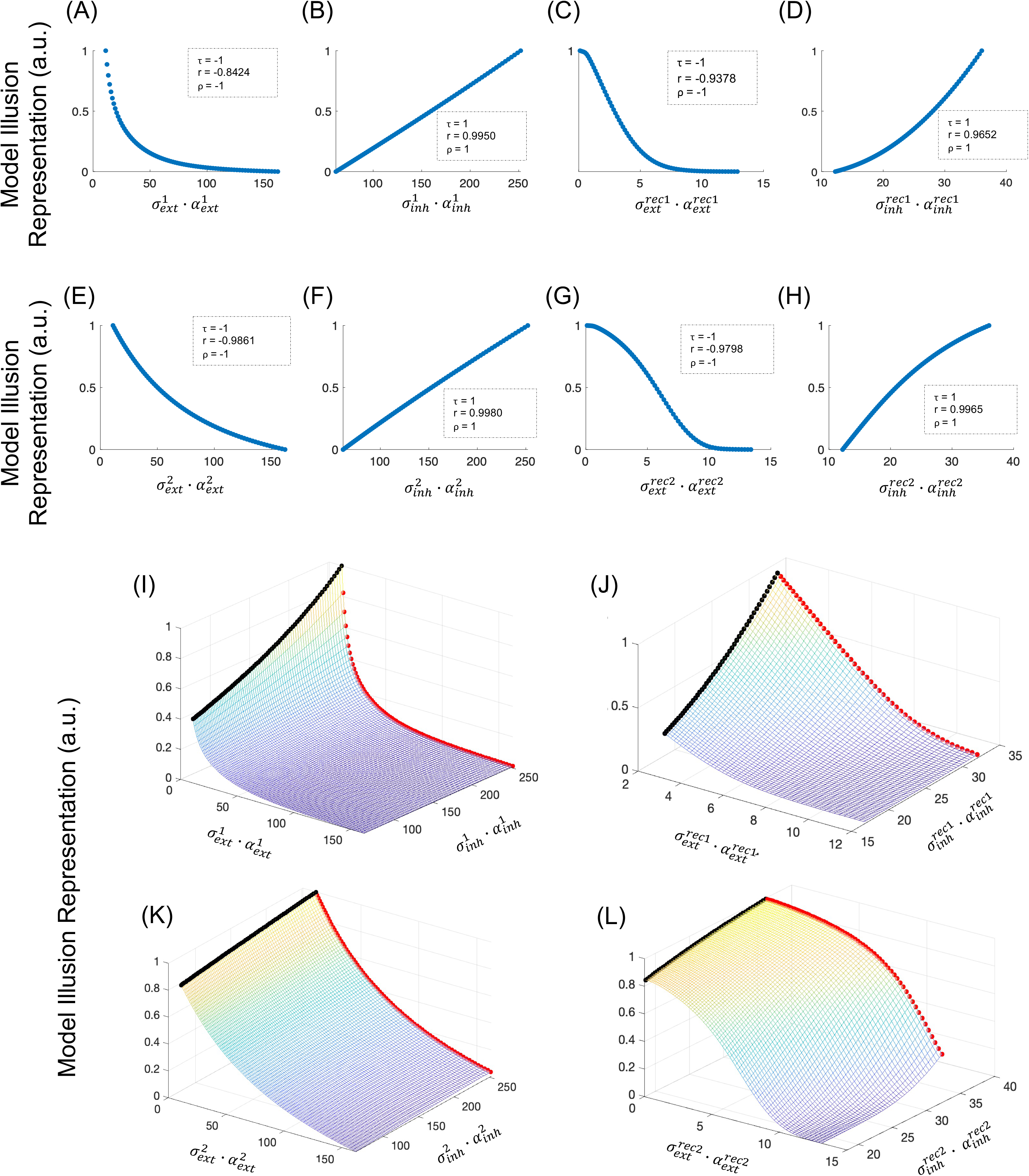
The effects of E/I modifications in the first and second layers, and recurrent connections of the neural model on Motion-Induced Position Shift (MIPS) illusion representation. MIPS illusion model representation against first-layer and second-layer excitatory subfield power (A/E), against inhibitory subfield power (B/F), against excitatory subfield power in recurrence (C/G) and against inhibitory subfield power in recurrence (D/H). (A/E) and (C/G) show that in a neural model, increasing the power of excitation in the first/second layers and their recurrent connections decrease the illusion representation; (B/F) and (D/H) show that increasing the power of inhibition in the first/second layer and their recurrent connections increase the amount of illusion representation. All correlation analysis show significance (see *p* values in Supplementary Table 1). Subplots (I-L) show that overall, in the neural model having feedforward, feedback, and recurrence, when excitatory subfield power in first and second layers, as well as their recurrent connections increase, the amount of illusion representation decreases (red); when inhibitory subfield power in first and second layers, as well as their recurrent connections increase, the amount of illusion representation increases (black). (I) and (J) demonstrate that the first layer is more sensitive to excitatory and inhibitory power modification, evidenced by the rapid decrease of high illusion representation (yellow) featured in neurotypical individuals. In comparison, (K) and (L) demonstrate that the second layer is more resistant to excitatory and inhibitory power modification, shown with larger persisting areas of high illusion representation (yellow) featured in neurotypical individuals. In addition, recurrent connections are more resistant to excitatory and inhibitory power modification compared to Inter-areal connections, shown by larger persisting area of high illusion representation (yellow) when recurrent subfield powers are modified (i.e. J compared to I, L compared to K).

To specifically investigate how the E and I subfield powers concurrently affect model illusion representation, we explored a wide range of E and I subfield power combinations (E/I ratio) in the surface plots of Figure 4I–L. These subplots show that in regions where the E power was strong and I power was weak (high E/I ratio), the model illusion representation was low (purple region). On the contrary, in regions where the E power was weak and I power was strong (low E/I ratio), the model illusion representation was high (yellow region).

Another insight drawn from the surface plots of Figure 4I–L is that the model illusion representation was less sensitive against E and I subfield power changes in layer 2 compared to layer 1. For example, comparing layer 1 and layer 2 inter-areal connectivity, the region of high illusion representation (yellow) characterized in neurotypical individuals showed a rapid decrease in Figure 4I but persisted over a larger area in Figure 4K. A more apparent difference was observed when comparing layer 1 and layer 2 intra-areal connectivity in Figure 4J with Figure 4L, where the illusion representation in layer 1 was more sensitive to E or I intra-areal connectivity modification than the layer 2 intra-areal connectivity. The region of neurotypical base-level illusion representation (yellow) only persisted for approximately 1/10 of the total area in Figure 4J and faded rapidly; on the other hand, the high illusion representation region persisted over half of the total area in Figure 4L.

The difference in illusion sensitivity to subfield power changes between layers 1 and 2 can also be observed in the plots of Figure 4A-H, such as when comparing the concavity of the illusion representation vs. inhibitory-power plot of layer 1(Figure 4D) and layer 2 (Figure 4H). Layer 1 plot in Figure 4D is concave up and monotonically increasing, indicating that the illusion representation was minimal when the I subfield power value was small, and only when the I subfield power value fully increased, the illusion representation could return to a neurotypical base amount. In contrast, the layer 2 plot in Figure 4H is concave down and monotonically increasing, indicating that even when the I subfield power became smaller, the model illusion representation sustained longer at the neurotypical base level in layer 2. Such evidence supported that the illusion representation was less sensitive to modifications in layer 2 parameters compared to layer 1.

In addition, illusion representation was less sensitive to E and I subfield power changes in intra-areal connectivity (recurrence) compared to inter-areal connectivity (feedforward/feedback), as shown in Figure 4K and 4L. Specifically, the region of neurotypical base-level illusion representation (yellow) showed a rapid decrease as soon as the E/I ratio increased in Figure 4K for inter-areal connectivity but persisted over a larger area in Figure 4L for the intra-areal connectivity plot. Approximately, the region of neurotypical base-level illusion representation (yellow) for inter-areal connectivity only persisted for 1/5 of the total area in Figure 4K and faded rapidly; on the other hand, the base-level illusion representation region for intra-areal connectivity persisted over half of the total area in Figure 4L. Such evidence indicated that the illusion representation was less sensitive to modifications in intra-areal connectivity than inter-areal connectivity.

The difference in illusion sensitivity to subfield power changes between intra-areal and inter-areal connectivity can also be observed in the plots of Figure 4A-H, such as when comparing the concavity of the illusion representation vs. the excitatory-power plot of inter-areal connectivity (Figure 4A&E) and intra-areal connectivity (Figure 4C&G). The shapes of Figure 4A and 4E were concave up and monotonically decreasing, with the model illusion representation having a steep drop as soon as the E subfield power value began to increase. In contrast, Figure 4C and 4G were concave down and decreasing when the E subfield power began to increase. Then, the shape reached an inflection point and became concave up and decreasing when the excitatory subfield power continued to increase. Such evidence indicated that the illusion representation was less sensitive to intra-areal connectivity than inter-areal connectivity.

### 3.2 The Effects of E and I Subfield Power in Altering Motion Repulsion (AMR)

Similarly, we explored the model illusion representation of altered motion repulsion (AMR) to gain further insight into the effect of individual E and I subfield power change as well as a more comprehensive variety of subfield power combination changes. We first established a baseline with a specific projection extent (*σ*) and projection strength (α) as the control combination, where there was the possibility to modify *σ* and α in both increasing and decreasing directions (Table 2). The results of the E/I ratio change combinations were grouped in three different ways: (1) overall E/I ratio deviation from the control baseline, irrespective of the individual constituents of I and E subfield power changes (Figure 5A); (2) I subfield power deviation from the control baseline, irrespective of the individual E subfield power change (Figure 5B); (3) E subfield power deviation from the control baseline, irrespective of the individual I subfield power change (Figure 5C). In addition, three subplots containing standard error of means are presented (Figure 5D corresponds to 5A, Fig. 5E to 5B, Fig. 5F to 5C). All 10 combinations of E/I ratio changes are also presented individually with a comprehensive subfield power profile to showcase the complexity of individual subfield power changes behind the overall E/I ratio change (Figure 5G)

**Figure 5.**
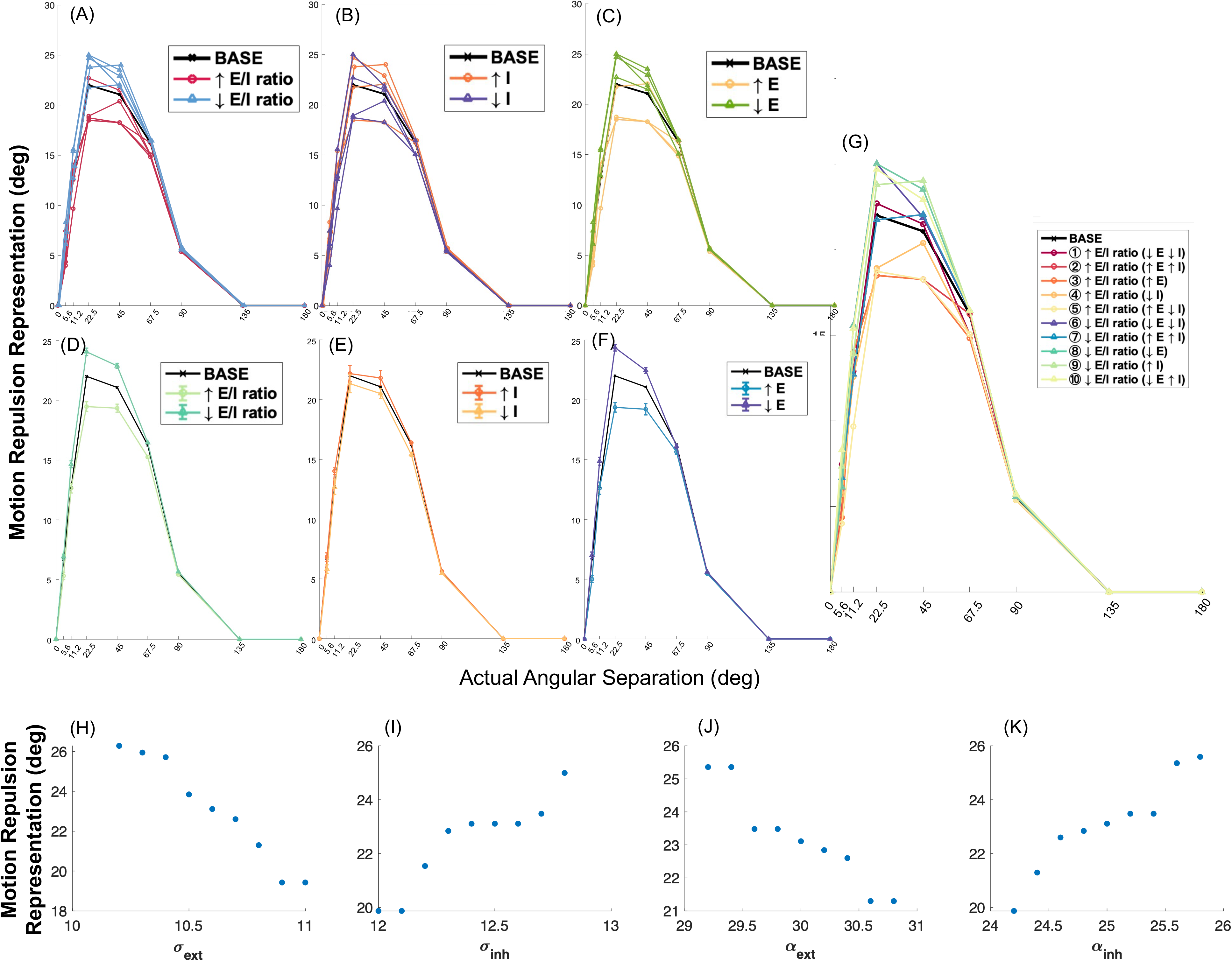
The effects of E/I on the model representation of motion repulsion illusion. The motion repulsion representation of the model depends on actual angular separation, with a range of 0° to 180°. BASE is a standard model with *σ_ext_*= 10.6, *σ_in_*_ℎ_= 12.4, *α_ext_*= 30.0, and *α_in_*_ℎ_= 25.0. (A) Compares the amount of motion repulsion representation of 5 ↑E/I combinations (red) and 5 ↓E/I combinations (blue) with respect to the BASE. (B) Compares the amount of motion repulsion representation of the 4 ↑I (orange) and 4 ↓I (purple) with respect to the BASE, and (C) compares the amount of motion repulsion representation of 4 ↑E (yellow) and 4 ↓E (green) with respect to the BASE. (D) Compares all ↑E/I combinations (dark green) and all ↓E/I combinations (light green) with respect to the BASE. (E) Compares all ↑I combinations (dark orange) and all ↓I combinations (light orange) with respect to the BASE. (F) Compares all ↑E combinations (blue) and all ↓E combinations (purple) with respect to the BASE. (G) Provides the details of all 10 combinations with the corresponding E/I ratio, E component changes, and I component changes, with respect to BASE. (H-K) show the Altered Motion Repulsion (AMR) illusion representation of the neural model when network excitatory and inhibitory subfields are modified. AMR illusion model representation against excitatory subfield extent (H), against inhibitory subfield extent (I), against excitatory subfield strength (J), against inhibitory subfield strength (K). (H) and (J) show that in the neural model, increasing the excitatory power, i.e. extent, strength, or both, decreases the illusion representation; (I) and (K) show that increasing the inhibitory power, i.e. extent, strength, or both, increases the amount of illusion representation. All correlation analysis show significance (see p values in Table 3).

**Table 3:** Mean and Standard Error of Mean (SEM) for AMR illusion representation.

|  | Mean for 22.5° | SEM for 22.5° | Mean for 45° | SEM for 45° |
| --- | --- | --- | --- | --- |
| Baseline* | 22.00 |  | 21.07 |  |
| ↑ E/I ratio | 19.47 | 0.4059 | 19.33 | 0.3393 |
| ↓ E/I ratio | 24.05 | 0.3095 | 22.88 | 0.2067 |
| ↑I | 22.18 | 0.6879 | 21.81 | 0.6253 |
| ↓I | 21.34 | 0.7641 | 20.50 | 0.4054 |
| ↑E | 19.37 | 0.3974 | 19.21 | 0.4733 |
| ↓E | 24.36 | 0.2788 | 22.45 | 0.2335 |

\*22.00 and 21.07 Baseline values are the Altered Motion Repulsion (AMR) illusion representation values at 22.5° and 45° instead of mean values

The angle between the reference direction and the test direction of dot movement is known as the actual angular separation in degrees, with values of 0°, 5.6°, 11.2°, 22.5°, 45°, 67.5°, 90°, 135°, and 180° (see methodology). For motion repulsion representation comparison, we focused on the *actual* angular separations of 5.6°, 11.2°, 22.5°, and 45°, based on findings in the literature ^28^. Notably, many of these angular separations demonstrated the most pronounced differences in *represented* angular motion separation compared to actual (motion repulsion representation) in our results, particularly at 22.5° and 45° (Figure 5A–G). The baseline combination showed peak motion repulsion at 22.5°. Conversely, when the angular separation exceeded 45°, the differences in motion repulsion representation became less pronounced.

In Figure 5A, we categorized the 10 combinations of E and I subfield changes into increased E/I ratio (red) and decreased E/I ratio (blue), irrespective of the individual constituents of I and E subfield power change. Compared with the control baseline combination (black line), most combinations with decreased E/I ratio had larger motion repulsion illusion representation at the actual angular separation of 22.5° and 45°. This was supported by the Mean ± SEM calculation at 22.5° and 45°, where the decreased E/I ratio had motion repulsion of 24.05 ± 0.3095 and 22.88 ± 0.2067 degrees respectively, greater than the 22.00 and 21.07 degrees respectively of the control baseline (Figure 5D and Table 3). On the other hand, most combinations with increased E/I ratio had smaller motion repulsion representations at 22.5° and 45° compared to the baseline. This was supported by the Mean ± SEM calculation of increased E/I ratio at 22.5° and 45° with results of 19.47 ± 0.4059 and 19.33 ± 0.3393 degrees respectively, which were less than the motion repulsion representation of 22.00 and 21.07 degrees respectively at the baseline (Figure 5D and Table 3). In general, the 5 combinations with increased E/I ratio demonstrated smaller illusion representation when directly compared to the 5 combinations with decreased E/I ratio, especially when comparing the illusion representation at the angular separation of 22.5° and 45°. The Mean ± SEM of increased E/I ratio was 19.47 ± 0.4059 and 19.33 ± 0.3393 degrees for 22.5° and 45°, smaller than 24.05 ± 0.3095 and 22.88 ± 0.2067 degrees respectively for decreased E/I ratio (Figure 5D and Table 3). Overall, these results suggested that an increase in E/I ratio is more representative of illusion susceptibility in AD with lower motion repulsion representation whereas a decrease in E/I ratio is more representative of neurotypical combinations with higher motion repulsion representation.

In Figure 5B, we categorized the 8 out of 10 combinations of E and I subfield changes into increased I subfield power (orange) and decreased I subfield power (purple), irrespective of the individual E subfield power change (the 2 combinations where only E subfield increased or decreased were excluded since they had no I subfield power changes and thus irrelevant). When comparing with the baseline combination (black line), an increased I subfield power had an overall larger motion repulsion representation at the actual angular separation of 45°. This was supported by the Mean ± SEM calculation at 45°, where increased I subfield power had motion repulsion of 21.81 ± 0.6253 degrees, greater than the 21.07 degrees value of the baseline (Figure 5E and Table 3). On the other hand, a decreased I subfield power had an overall smaller motion repulsion representation at the actual angular separation of 45°. This was supported by the Mean ± SEM calculation of decreased I subfield power at 45° with results of 20.50 ± 0.4054 degrees, which were less than the motion repulsion representation of 21.07 degrees at the baseline (Figure 5E and Table 3). In general, the 4 combinations with increased I subfield power demonstrated larger motion repulsion representation compared to the 4 combinations with decreased I subfield power, when comparing the motion repulsion at angular separation of 45°. The Mean ± SEM of increased I subfield power was 21.81 ± 0.6253 for 45°, larger than 20.50 ± 0.4054 degrees for decreased I subfield power (Figure 5E and Table 3). Overall, these results suggested that decreased I subfield power is more representative of illusion susceptibility in AD with lower motion repulsion representation whereas increased I subfield power is more representative of neurotypical combinations with higher motion repulsion representation.

In Figure 5C, we categorized the 8 out of 10 combinations of E and I subfield changes into increased E subfield power (yellow) and decreased E subfield power (green), irrespective of the individual I subfield power change (the 2 combinations where only the I subfield increased or decreased were excluded since they had no E subfield power changes and thus irrelevant). When comparing with the baseline combination (black line), most combinations with an increased E subfield power had smaller motion repulsion representation at the actual angular separation of 22.5° and 45°. Based on the Mean ± SEM calculation at 22.5° and 45°, increased E subfield power had a motion repulsion of 19.37 ± 0.3974 and 19.21 ± 0.4733 degrees respectively, less than the 22.00 and 21.07 degrees respectively of the baseline (Figure 5F and Table 3). On the other hand, all combinations with a decreased E subfield power had larger motion repulsion representation at the actual angular separation of 22.5° and 45° when compared to the baseline. This was supported by the Mean ± SEM calculation of decreased E subfield power with a result of 24.36 ± 0.2788 and 22.45 ± 0.2335 degrees respectively, more than the 22.00 and 21.07 degrees respectively of the baseline (Figure 5F and Table 3). In general, the 4 combinations with increased E subfield power demonstrated smaller motion repulsion representation compared to the 4 combinations with decreased E subfield power, especially at an angular separation of 22.5° and 45°. The Mean ± SEM of increased E subfield power was 19.37 ± 0.3974 and 19.21 ± 0.4733 degrees for 22.5° and 45° respectively, smaller than 24.36 ± 0.2788 and 22.45 ± 0.2335 degrees of decreased E subfield power (Figure 5F and Table 3). These results suggested that increased E subfield power is more representative of illusion susceptibility in AD with lower motion repulsion representation whereas decreased E subfield power is more representative of neurotypical combinations with higher motion repulsion representation.

Overall, the motion repulsion representation reached peak levels at an angular separation of 22.5° or 45° for all combinations and decreased for an angular separation of 45° and onward. The two combinations with the largest motion repulsion representation representative of neurotypical combination were: 1) decreased E/I ratio and decreased E subfield power with average illusion representation of 24.27° degree at the actual angular separation of 22.5° or 45° (⑧ in Figure 5G), 2) decreased E/I ratio and increased I subfield power with average illusion representation of 23.91° (⑨ in Figure 5G). The two combinations with the smallest illusion representation representative of AD combination at the actual angular separation of 22.5° or 45° were: 1) increased E/I ratio and increased E subfield power and increase I subfield power (② in Figure 5G), 2) increased E/I ratio, increased E subfield power (③ in Figure 5G), with average motion repulsion illusion representation of 18.38° degree.

Interestingly, our results also suggested stronger influence of E part than I part in the overall E/I ratio, aligning with the idea that Aβ-induced synaptic dysfunction primarily affects excitatory synapses^7,15,50^. So, our findings suggest that there is more promising potential to place an emphasis on future treatment strategies focused on modulating excitatory circuits. The stronger E influence was demonstrated by the cases of exceptions to general trends in the results. One exception to the general trend of increased E/I ratio leading to decreased illusion representation occurred when the overall E/I ratio increased while both E and I subfield power decreased (① in Figure 5G). In this case, the combination resulted in slightly higher illusion representation than the baseline. This could indicate some dominating influence from E subfield power, the decreased E subfield power helped retain higher illusion representation than baseline despite the overall increased E/I ratio. Another case of stronger E influence was an exception to the general trend of increased I subfield power leading to higher illusion representation. When both the E and I subfield power increased along with the overall E/I ratio (② in Figure 5G), despite the increase in I power, the combination resulted in a reduced illusion representation compared to the baseline. This reiterated some dominating influence from E subfield power, the increased E subfield power reduced illusion representation compared to baseline, nullifying the effect of increased I subfield power.

To reflect the intricacy behind the overall E/I ratio change and break down the effects of changing individual E and I power constituents on motion repulsion illusion representation, the results of different modifications in the model parameter constituents (*σ_ext_*, *σ_in_*_ℎ_*, α_ext_*, and *α_in_*_ℎ_) of subfield powers are shown (Figure 5H–K). Only individual constituents were changed one at a time, keeping the other parameters at baseline values to offer more insights about the contribution from each constituent. Overall, the results indicated that when *σ_ext_* of inter-areal and intra-areal connectivity in layer 1 and 2 increased, the amount of motion repulsion illusion representation decreased (Figure 5H). When *α_ext_* increased, the amount of illusion representation also decreased (Figure 5J). On the other hand, when *σ_in_*_ℎ_ increased, the amount of illusion representation increased (Figure 5I). When *α_in_*_ℎ_ increased, the amount of illusion representation also increased (Figure 5K). Therefore, taking into account the consistent trends of *σ* and *α* shown, we can summarize that when there is an overall increase in the E/I ratio, (*σ_ext_* × *α_ext_*)/(*σ_in_*_ℎ_ × *α_in_*_ℎ_), the amount of motion repulsion representation decreases in AMR.

## 4. Discussion

This study, developed a two-layer rate based neural model, representing the early and higher visual areas with feedforward, feedback, and recurrent connections. We used Motion-Induced Position Shift (MIPS) and Altered Motion Repulsion (AMR) illusions as inputs, to explore the relationship between excitatory and inhibitory (E/I) neural connectivities and their influence on visual motion perception in Alzheimer’s Disease (AD). The alteration in visual processing, including motion perception, is one of the early cognitive signs of AD, often preceding memory impairment and other cognitive deficits. Studies have demonstrated that individuals with AD experience difficulties in various visual tasks, such as pattern recognition, spatial orientation, and motion detection ^51–53^. These deficits are closely linked to structural and functional changes in brain regions that process visual information, such as the primary visual cortex (V1) and higher-order visual areas ^54^. By factorial assessment and analyzing the neural model representation of visual motion illusions, our study revealed 4 key insights: 1) When excitatory (E) subfield power increased, the amount of illusion representation decreased; when inhibitory (I) subfield power decreased, the amount of illusion representation decreased. These two conditions were more representative of reduced illusion susceptibility in AD. On the other hand, when excitatory (E) subfield power decreased, the amount of illusion representation increased; when inhibitory (I) subfield power increased, the amount of illusion representation increased. These two conditions were more representative of neurotypical conditions. 2) When considering E and I subfield power concurrently, an increased E/I ratio from the base condition was more representative of reduced illusion susceptibility in AD. 3) Model illusion representation in model layer 2 (representing higher visual processing) was more resistant against E and I subfield power changes compared to layer 1 (representing lower visual processing). 4) Model illusion representation demonstrated less sensitivity to reduction when the E/I ratio of intra-areal connectivity (recurrence) increased, while the illusion representation demonstrated more sensitivity to reduction when the E/I ratio of inter-areal connectivity (feedforward and feedback) increased. These four findings provide a framework for investigating how specific alterations in excitatory versus inhibitory components, different visual processing stages, and intra-areal versus inter-areal connectivity in AD may influence changes in visual illusion susceptibility, offering insights into the neural mechanisms underlying visual processing deficits in AD.

### 4.1 Impact of Individual Excitation or Inhibition Changes on Neuronal dynamics and Pathophysiology

Excitatory neurons are related to AD pathology mainly through glutamate-induced excitotoxicity ^9,55–57^. Glutamate signaling is normally highly regulated. However, in brains with neurodegeneration, impaired glutamate clearance leads to overactivation of NMDA receptors, triggering intracellular cascades, resulting in neuronal death ^58,59^. This excitotoxicity likely contributes to neuronal loss and synaptic changes observed in AD ^10,60,61^. Our findings showed increased excitatory power correlates with reduced susceptibility to visual motion illusions, suggesting that hyperexcitation alters perception, further supporting the excitotoxicity hypothesis.

Aβ-induced synaptic dysfunction primarily affects excitatory synapses, reducing synaptic strength and leading to connection loss ^62–64^. Moreover, hyperphosphorylated Tau in AD and other neurodegenerative disorders is at least partially trans-synaptically transmitted through cortical pathways, likely affecting synaptic function ^65,66^. Compensatory hyperactivity, impaired glutamate clearance, and toxic extrasynaptic receptor activation amplify excitotoxic conditions despite weakened synaptic activity, explaining a paradoxical coexistence of synaptic loss and hyperexcitation. Excitatory neurons in AD also exhibit disrupted calcium homeostasis, where NMDA overactivation leads to calcium overload, mitochondrial dysfunction, oxidative stress, and cell death ^67,68^. Our study suggested that such hyperactivity may disrupt visual processing, reflecting broader effects of Aβ and neurofibrillary tangles on circuit function.

Inhibitory signaling is also altered in AD. GABA, the primary inhibitory neurotransmitter, maintains circuit balance, and its dysregulation in AD is well-documented ^11,16–19^. Degeneration of GABAergic interneurons, such as fast spiking, parvalbumin-positive (PV+) neurons, reduces cortical inhibition and impairs cognitive function. Such loss disrupts neural synchrony and oscillation, contributing to cognitive abnormalities ^69^. Our study showed that decreased inhibitory (I) subfield power correlated with reduced motion illusion representation, suggesting that impaired inhibition may underlie perceptual deficits in AD. Enhancing inhibition may therefore offer palliative and therapeutic value.

### 4.2 Impact of Excitation/Inhibition Ratio Changes on Neuronal dynamics and Pathophysiology

Although excitation and inhibition exert distinct effects, the excitatory/inhibitory (E/I) ratio offers a more holistic view of circuit function. This ratio is critical for circuit stability and sensory processing, and its disruption is linked to various neuropsychiatric and neurodegenerative disorders ^14,70^. The E/I ratio in AD reflects glutamate-induced hyperexcitation and impaired inhibition ^18,56,58,69^. Together, these changes disrupt the E/I balance, creating network dysfunctions. Our study supports this, showing increased E/I ratio in visual areas correlates with reduced motion illusion susceptibility. Conversely, strengthened inhibition (lower E/I ratio) enhances illusion perception. These results highlight E/I ratio as a meaningful indicator of neural imbalance in sensory circuits like V1 and MT/V5. By framing the E/I ratio as a standalone measure, distinct from isolated excitatory or inhibitory contributions, our study emphasized its relevance in assessing neural dysfunction. Future research should explore how E/I ratio can relate to specific AD symptoms, incorporating physiological and computational approaches.

### 4.3 Impact of Changes in Early Sensory Cortices vs. Higher Cortices on Neuronal Dynamics and Pathophysiology

AD affects cortical regions differently. Higher-order areas (e.g. in the temporal lobes) show earlier changes than early sensory regions (e.g. V1), but higher-order may decline more slowly due to structural resilience ^71–74^. Age-related declines in plasticity and repair mechanisms may contribute to higher-order changes early on ^75^. Our results supported this difference: layer 2 (modeling higher-order areas) maintained visual motion illusion representation better than layer 1 (early sensory), suggesting higher-order areas might be more resilient against E/I imbalances. In contrast, early areas like V1, which require precise E/I balance may be more sensitive to disruptions. These findings underscored the importance of studying E/I dynamics across sensory hierarchy to understand AD progression.

### 4.4 Impact of Recurrence vs. Feedforward and Feedback Connections on Neuronal Dynamics and Pathophysiology

Our findings revealed that neurodegeneration affects long-range and short-range/local connectivity differently. Long-range connectivity via white matter facilitates global communication but can be highly vulnerable in early AD, correlating with cognitive decline ^76–79^. Short-range local circuits exhibit initial resilience and may enhance to compensate, supporting recurrent neural processing ^80,81^. Nevertheless, as AD progresses, grey matter deterioration impairs even local processing, nullifying compensation ^82,83^. Feedforward, feedback, and recurrent connections are all disrupted: long-range loss impairs global integration, while short-range circuits initially compensate then fail. Interventions can adopt these strategies: preserving white matter could maintain long-range connections and fortifying local circuits early on.

### 4.5 Early Intervention and Cognitive Assessment Tools

Our study highlighted the potential of visual illusions like MIPS and AMR to supplement existing diagnostic methods for a non-invasive and cost-effective way of early AD detection since perceptual deficits emerge early. Early diagnosis allows intervention that can slow disease progression and improve life quality ^84^. Currently, biomarkers (Aβ, tau) often appear after significant loss, lowering intervention efficacy ^85^. Our findings suggested E/I imbalance and perceptual alterations could serve as earlier indicators. Besides AD, our modeling platform could extend to other disorders with known E/I imbalance: Schizophrenia, ASD, Parkinson’s disease, or epilepsy ^86–89^.

### 4.6 Limitations

1) Quantification Methods for Representation of Visual Illusions: While we proposed potential readout quantifications for model representations of MIPS and AMR, empirical work is needed to confirm how the brain represents such illusions. Our current metrics were based on information availability of neural signals, but how they are read out, demands further empirical investigation. 2) Additional Neurobiological Parameters: We considered E, I, and the E/I ratio as neurobiological parameters influencing our model’s representation of illusions, but other neurobiological parameters such as dendritic integration (that is represented in our model by “extent”:σ), synaptic plasticity, or modulatory systems (cholinergic, dopaminergic) could also influence E/I balance ^90–92^, and therefore, our future direction would be to explicitly characterize the impact of such neurobiological parameters on the current study’s αs and σs. Characterizing such neurobiological factors that affect E/I, together with how altered E/I changes model visual representation, will enable a more detailed and biologically informed approach. 3) Exploration of Other Visual Illusions: Currently, AD presents a broad spectrum of visual symptoms ranging from lower-level deficits to higher-level impairments ^20–26^. Yet, there is limited literature on specific visual illusions such as MIPS and AMR. While our focus on MIPS and AMR was supported by psychometric studies and their correlation with MMSE scores, there is a clear need for further psychophysical studies exploring changes in other visual illusions in AD. Visual phenomena such as simultaneous contrast, contrast-contrast illusions, and apparent motion illusions, which have been investigated in Autism Spectrum Disorder and Schizophrenia, provide valuable insights into the neurobiological changes underlying these conditions ^34–36^. Extending this research to AD could deepen our understanding of perceptual and neurocircuitry alterations. 4) Expansion of Computational and Neurophysiological Paradigm Scope: Although our study directly compared the model’s predictions with existing psychophysical data related to illusion perception in both healthy individuals and those with AD, future scope expansions could explore additional cortical areas within the visual pathway beyond the early visual and higher order cortices represented by two layers currently modeled or even other cognitive modalities such as auditory processing, which is significantly impacted in AD ^93^. Further validation using physiological and imaging methods would be critical for enhancing both the biological relevance and the translational potential of our findings.

## Abbreviations

AD: Alzheimer’s disease;
E/I: excitatory/inhibitory;
MIPS: Motion-Induced Position Shift;
AMR: Altered Motion Repulsion;
V1: primary visual cortex;
MT/V5: middle temporal area/visual area 5.

## Data Availability Statement

The code used to support the findings of this study has been deposited in a public repository and is openly accessible at https://github.com/BUicicchen/Alzheimers-Computation-Research

## Consent Statement

Consent was not necessary: this study is a computational modeling work based on published and public data, no human subjects were recruited for this study and as such, consent was not necessary.

---

**Supplementary Table 1:**
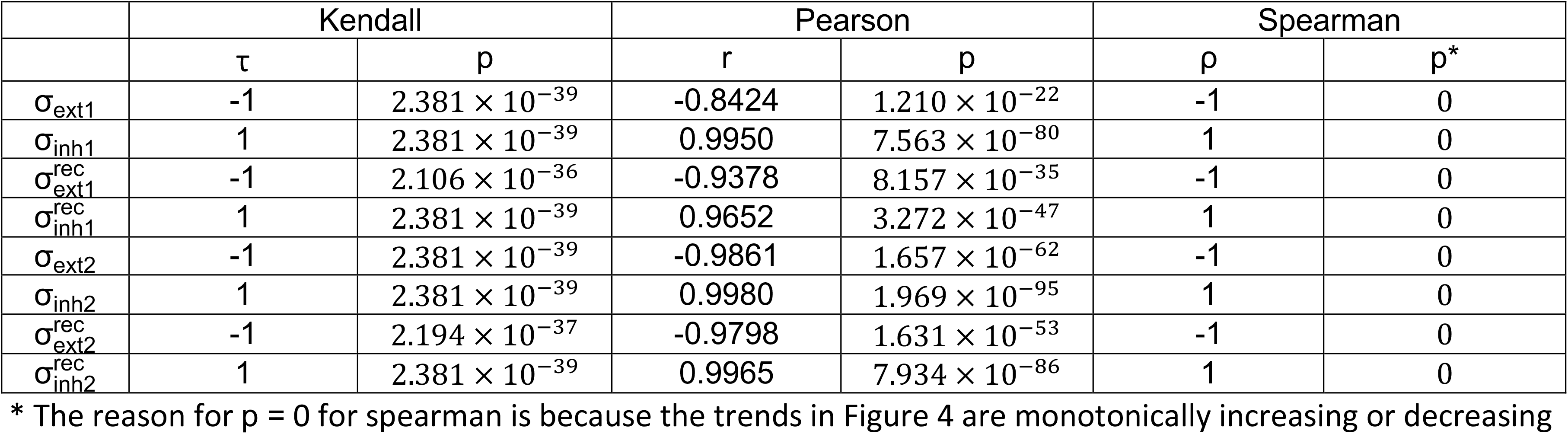
Statistical measures of correlation for MIPS illusion representation.

|  | Kendall |  | Pearson |  | Spearman |  |
| --- | --- | --- | --- | --- | --- | --- |
| | $\tau$ | p | r | p | $\rho$ | p* |
| $\sigma_{\text{ext}1}$ | -1 | $2.381 \times 10^{-39}$ | -0.8424 | $1.210 \times 10^{-22}$ | -1 | 0 |
| $\sigma_{\text{inh}1}$ | 1 | $2.381 \times 10^{-39}$ | 0.9950 | $7.563 \times 10^{-80}$ | 1 | 0 |
| $\sigma_{\text{ext}1}^{\text{rec}}$ | -1 | $2.106 \times 10^{-36}$ | -0.9378 | $8.157 \times 10^{-35}$ | -1 | 0 |
| $\sigma_{\text{inh}1}^{\text{rec}}$ | 1 | $2.381 \times 10^{-39}$ | 0.9652 | $3.272 \times 10^{-47}$ | 1 | 0 |
| $\sigma_{\text{ext}2}$ | -1 | $2.381 \times 10^{-39}$ | -0.9861 | $1.657 \times 10^{-62}$ | -1 | 0 |
| $\sigma_{\text{inh}2}$ | 1 | $2.381 \times 10^{-39}$ | 0.9980 | $1.969 \times 10^{-95}$ | 1 | 0 |
| $\sigma_{\text{ext}2}^{\text{rec}}$ | -1 | $2.194 \times 10^{-37}$ | -0.9798 | $1.631 \times 10^{-53}$ | -1 | 0 |
| $\sigma_{\text{inh}2}^{\text{rec}}$ | 1 | $2.381 \times 10^{-39}$ | 0.9965 | $7.934 \times 10^{-86}$ | 1 | 0 |

\* The reason for p = 0 for spearman is because the trends in Figure 4 are monotonically increasing or decreasing

